# O-GlcNAc transferase suppresses necroptosis and liver fibrosis

**DOI:** 10.1101/519975

**Authors:** Bichen Zhang, Min-Dian Li, Ruonan Yin, Yuyang Liu, Yunfan Yang, Kisha A. Mitchell – Richards, Jin Hyun Nam, Rui Li, Li Wang, Yasuko Iwakiri, Dongjun Chung, Marie E. Robert, Barbara E. Ehrlich, Anton M. Bennett, Jun Yu, Michael H. Nathanson, Xiaoyong Yang

## Abstract

Over a billion people suffer from chronic liver diseases worldwide, which often leads to fibrosis and then cirrhosis. Treatments for fibrosis remain experimental, in part because no unifying mechanism has been identified that initiates liver fibrosis. Here we report that O-linked β-N-acetylglucosamine (O-GlcNAc) modification protects against hepatocyte necroptosis and initiation of liver fibrosis. Decreased O-GlcNAc levels were seen in patients with liver cirrhosis and in mice with ethanol-induced liver injury. Liver-specific O-GlcNAc transferase (OGT) knockout (OGT-LKO) mice exhibited ballooning degeneration and elevated circulating alanine aminotransferase (ALT) levels at an early age and progressed to liver fibrosis and portal inflammation by 10 weeks of age. OGT-deficient hepatocytes underwent excessive necroptosis and exhibited elevated protein expression levels of receptor-interacting protein kinase 3 (RIPK3) and mixed lineage kinase domain-like (MLKL), which are key mediators of necroptosis. Furthermore, glycosylation of RIPK3 by OGT reduced RIPK3 protein stability. Taken together, these findings identify OGT as a key suppressor of hepatocyte necroptosis and OGT-LKO mice may serve as an effective spontaneous genetic model of liver fibrosis.

## Introduction

More than a billion people worldwide are afflicted by chronic liver disease, the most common causes of which include fatty liver, infection with Hepatitis B or C virus, and alcohol. Despite differences in the etiology of these chronic liver diseases, they each lead to excessive stress on the liver and result in hepatocyte death(1). Hepatocellular death contributes significantly to chronic liver disease by inducing inflammatory responses that lead to hepatic fibrosis and cirrhosis (2). Recent evidence suggests that necroptosis, a form of caspase-independent, regulated cell death, is manifested in a variety of acute and chronic types of liver diseases(3, 4). Necroptosis has been implicated in patients with alcoholic steatohepatitis (ASH), non-alcoholic steatohepatitis (NASH), and hepatitis B or C infections(5, 6), underscoring its prevalence and general role in the pathogenesis of chronic liver diseases.

Necroptosis is mediated by the “necrosome”, a signaling protein complex that is composed of receptor-interacting protein kinase 3 (RIPK3) and mixed lineage kinase domain-like (MLKL)(7). Phosphorylated RIPK3 recruits and activates MLKL, which translocates to the cell membrane and causes membrane rupture(8). Owing to the loss of membrane integrity in necroptotic cells, the intracellular contents act as damage-associated molecular patterns (DAMPs) that recruit immune cells and cause sterile inflammation in affected tissues(9). Necroptosis also promotes cell-autonomous production of pro-inflammatory cytokines(10). Therefore, necroptosis is considered as a highly inflammatory form of cell death. In light of the data collected from patients, a myriad of studies in recent years have investigated the contribution of necroptosis in different mouse liver injury models and demonstrated the importance and complexity of necroptosis in these pathological conditions (11–16). However, the mechanism that governs necroptotic hepatocyte death is largely unknown.

O-GlcNAcylation is a dynamic and reversible post-translational modification (PTM) that regulates a broad spectrum of cellular events ranging from epigenetics and transcription to signaling and rhythm(17–20). By sensing and integrating the nutrient and stress cues, it prepares cells to respond promptly to an ever-changing microenvironment. A pair of enzymes, O-GlcNAc transferase (OGT) and O-GlcNAcase (OGA), actively controls the cycling of O-GlcNAc modification on serine or threonine residues of cytoplasmic, nuclear, and mitochondrial proteins. Preserving the O-GlcNAc level in an optimal zone is essential for maintaining cellular homeostasis, whereas aberrant O-GlcNAcylation is linked to a plethora of diseases including cancer, metabolic syndromes, and cardiovascular diseases(21–23). Throughout the years it has become evident that dysfunction of O-GlcNAcylation sensitizes cells to various kinds of stress ultimately leading to cell death(24, 25). This phenomenon has been repeatedly reported in various tissues and pathologies(26–29), yet the mechanisms by which the perturbation of O-GlcNAcylation affects cell survival and death is undefined.

Previously, we have shown that OGT performs essential functions in liver physiology by regulating gluconeogenesis and the insulin signaling pathway(30, 31), circadian clock(32), and autophagy(33). Germline OGT deletion is lethal at the embryogenesis stage. Our previous models studying hepatic OGT are based on transient genetic interventions. A major caveat is that we did not know the chronic effects of OGT deletion on hepatocytes. Here, we report that chronic hepatocyte-specific deletion of OGT leads to a spontaneous, rapidly developing liver injury due to massive hepatocellular necroptosis. We further demonstrate that OGT serves as a negative regulator of necroptosis by suppressing RIPK3 expression. These findings reveal that hepatocyte OGT protects the liver against necroptosis and thus liver fibrosis.

## Materials and methods

### Mice and human liver samples

*Ogt^flox^* mice on C57BL/6J background were provided by Dr. Steven Jones at the University of Louisville (34). *Alb-Cre* mice were provided by Dr. Anton Bennett at Yale University. All mice were kept on a 12 hr light, 12 hr dark cycle. All mice have free access to food and water. Mice were fasted for 6 hr before sacrifice and tissues were collected for protein and RNA extraction and histology analysis. Ethanol-induced liver injury was induced as described before(35). Briefly, mice were fed with a Lieber-DeCarli liquid diet (F1258SP, BioServ, Flemington, NJ) or a control diet (F1259SP, BioServ) that has a matched calorie with ethanol diet for 6 weeks. All procedures have been approved by the Institutional Animal Care and Use Committee of Yale University. Both genders were included in the experiments and the age of mice used is specified in the text. The human liver samples were provided by Dr. Li Wang at the University of Connecticut. The samples were obtained from Liver Tissue Cell Distribution System (Minneapolis, Minnesota) as described before(36).

### Metabolic assays

Body weights were measured each week. Metabolic cage study was carried out with the Promethion multiplexed metabolic measurement system. Mice were single-housed for five days and acclimated in metabolic chambers for three days before the measurement of gas exchange, food intake, and ambulatory activity.

### Histology

Mouse livers were dissected and fixed in 4% paraformaldehyde for 48 hr. Hematoxylin and eosin stains and Masson’s trichrome stains were performed by the Histology Core in the Department of Comparative Medicine. Immunohistochemistry was performed with the Vectastain Elite ABC HRP kit (Vector Laboratories, Burlingame, CA). Paraffin-embedded sections were de-paraffined and antigen retrieval was carried out in a preheated steamer. The slides were then blocked for background peroxidase activity with hydrogen peroxide solution and unspecific binding with blocking buffer provided in the kit. The slides were incubated with 1:50 primary antibody overnight at 4°C and secondary antibody for 30 min at room temperature. The slides were counterstained with Meyer’s hematoxylin solution, dehydrated, and mounted.

### ELISA

Liver tissues were weighed and homogenized in assay diluent provided by the TNF-α or IL-6 ELISA kit (BD, Franklin Lakes, NJ). Plasma samples were collected with heparin as anticoagulant, centrifuged at 1000×g at 4°C, and stored at -80°C until measurement. Assays were performed according to the manufacturer’s protocol.

### RNA sequencing

mRNA was extracted from snap-frozen liver tissues with QIAGEN RNeasy (QIAGEN, Hilden, Germany) kit. Library preparation and sequencing were carried out at Novogene (Beijing, China) on the Illumina NovaSeq 6000 system with total raw reads at 20 million per sample on average. RNA sequencing data analysis was performed in collaboration with Dr. Dongjun Chung’s group at the Medical University of South Carolina.

### Flow cytometry

Freshly isolated primary hepatocytes were washed twice with BioLegend’s cell staining buffer and then resuspended in Annexin V binding buffer at the density of 5×10^6^ cells/ml. 100 μl cell suspension was transferred to the test tube and stained with 5 μl Pacific Blue Annexin V and propidium iodine (PI) solution (BioLegend, San Diego, CA) respectively for 15 min in dark. 400 μl Annexin V binding buffer was then added to the test tube and the samples were analyzed with BD LSR II flow cytometer.

### Fluorescent imaging and live cell imaging

Primary hepatocytes were plated on 8-well chamber slides for immunofluorescence imaging and on collagen-coated 6-well plates (BD, Franklin Lakes, NJ) for PI/Hoechst stains and MitoSox stains. For immunofluorescence, primary hepatocytes were fixed with 4% paraformaldehyde for 20 min and permeabilized in 0.5% Triton X-100 for 10 min at room temperature. Cells were blocked with 2% BSA in PBST (0.5% Tween 20 in PBS) and then incubated with primary antibodies at 4°C overnight in a humidified chamber. The cells were incubated with Alexa Fluor 488-conjugated secondary antibodies for 1 hr at room temperature in dark followed by staining with DAPI. The slides were mounted with Vectashield antifade mounting medium (Vector Laboratories, Burlingame, CA). For PI/Hoechst staining, primary hepatocytes were washed with PBS and incubated first with PI (BioLegend, San Diego, CA) and then with Hoechst 33342(Thermo Fisher, Waltham, MA) in dark at room temperature for 5 min. MitoSox Red (Thermo Fisher, Waltham, MA) staining was performed according to the manufacturer’s protocol.

### Cell viability assays

The CellTiter-Glo luminescent cell viability assay kit (Promega, Madison, WI) was used to determine the viability of primary hepatocytes. The cells were plated in black-wall, clear-bottom 96-well plate at the density of 3×10^4^ cells/well. 4 hr after plating, the cells were treated with various drugs as indicated in the text for 24 hr. The plates were brought to room temperature for 30 min equilibration and 100μl assay buffer was added to each well. The plates were incubated for 10 min and luminescence was determined with the Tecan Infinite M200 plate reader.

### Cell culture

HEK293T, H1299, and 3T3-L1 cells were cultured in DMEM with 10% fetal bovine serum (FBS). LX-2 cells were cultured in DMEM with 2% FBS. Primary hepatocytes from WT and OGT-LKO mice were isolated by Yale Liver Center Core Facility and cultured in Williams’ Medium E supplemented with 10% FBS, 10 mM HEPES buffer, 2 mM L-glutamine, 8 mg/L gentamicin, SPA, 1 μM dexamethasone, 4 μg/ml insulin, and 1 mM glucose. Human primary hepatocytes were provided by the Yale Liver Center Translational Core Facility and plated in HMM medium (Lonza, Basel, Switzerland) supplemented with 100 nM insulin, 100 nM dexamethasone, 50 μg/ml gentamicin, and 50μg/ml amphotericin. All primary hepatocytes were plated on Collagen I coated plates (BD, Franklin Lakes, NJ). Transfection of plasmids or siRNAs was performed with FuGene HD (Promega, Madison, WI) or Lipofectamine RNAimax (Invitrogen, Carlsbad, CA) respectively. DMEM, FBS, William’s medium E, HEPES buffer, glutamine stock solution, SPA, gentamicin, amphotericin were from Gibco (Waltham, MA). Dexamethasone, insulin, and glucose were purchased from Sigma-Aldrich (St. Louis, MO).

### Plasmids and siRNAs

Myc-OGT was kindly provided by Dr. Xiaochun Yu at the University of Michigan. The mRIPK3-GFP plasmid was purchased from Addgene. FLAG tag at the C-terminal of human RIPK3 was cloned into the pEGFP-N1 vector. Scrambled siRNA (5’-GAGGCAUGUCCGUUGAUUCGU-3’) and OGT siRNA (5’-GAGGCAGUUCGCUUGUAUCGU -3’) were synthesized by Dharmacon (Lafayette, CO).

### Real-time PCR

Total RNA was extracted from cells or tissues with TRIzol reagent (Invitrogen, Carlsbad, CA). Complementary DNA was synthesized from total RNA with Superscript III enzyme (Bio-Rad, Hercules, CA) and amplified with SYBR Green Supermix (Bio-Rad, Hercules, CA) using a LightCycler480 Real-Time PCR system (Roche, Basel, Switzerland). All data were normalized to the expression of 36B4. Primer sequences are listed in Table S2.

**Table S2.**
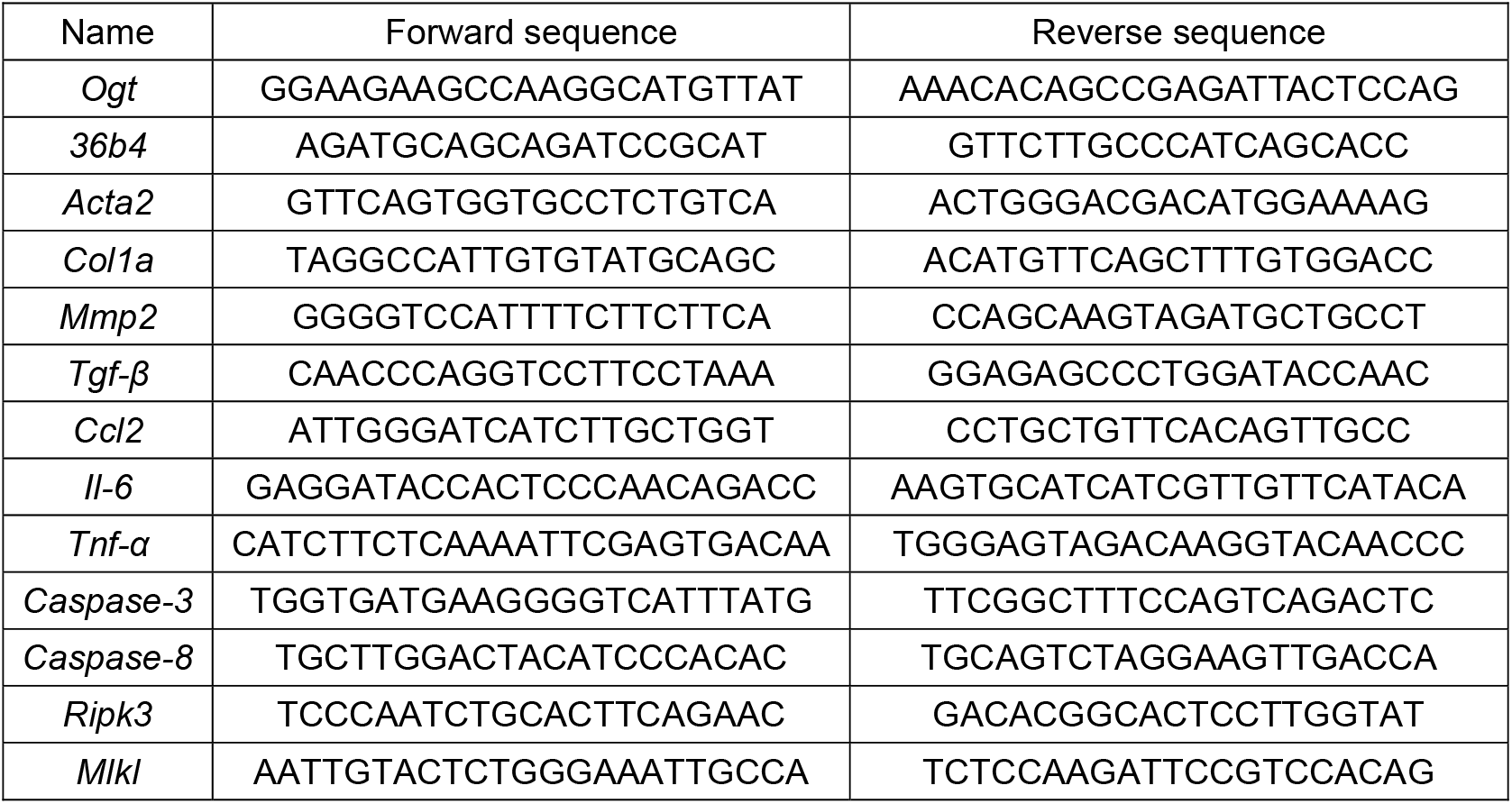
Primer sequences

### Western blotting and immunoprecipitation

Liver tissues or cells were lysed in buffer containing 1% Nonidet P-40, 150 mM NaCl, 0.1 mM EDTA, 50 mM Tris-HCl, proteinase inhibitors, and protein phosphatase inhibitors. In immunoprecipitation assays, cell lysates were mixed with various antibodies and precipitated with protein A/G agarose beads (Santa Cruz Biotechnology, Dallas, TX) or FLAG M2 monoclonal antibody affinity gel (Sigma-Aldrich, St. Louis, MO) overnight at 4°C. The lysates were then washed and boiled in SDS loading buffer. Equal amounts of protein lysates were resolved on SDS-PAGE gels and transferred to PVDF membrane. The membranes were blocked in 5% BSA and incubated with various primary antibodies overnight at 4°C. After three washes, the membranes were incubated with peroxidase-conjugated secondary antibodies for 1 hr and visualized with ECL chemiluminescent substrate.

### Antibodies and reagents

Antibodies against OGT (ab96718), O-GlcNAc (RL2, ab2739), p-MLKL S345 (ab196436) (Abcam, Cambridge, UK); cleaved caspase-3 (9664), cleaved caspase-8 (8592), RIPK1 (3493), FLAG (2368), β-tubulin (2128) (Cell Signaling Technology, Danvers, MA); Myc (sc-40), RIPK3 (sc-374639), β-actin (sc-8432) (Santa Cruz Biotechnology, Dallas, TX); F4/80 (14-4801-82, eBioscience, Waltham, MA) were purchased from the indicated sources. Antibodies against p-RIPK3 and MLKL were kindly provided by Dr. Vishva Dixit from Genentech (San Francisco, CA). Alexa Fluor 488-conjugated secondary antibodies were obtained from Thermo Fisher Scientific (Waltham, MA). DAPI, Hoechst 33342, Trypan blue solution, and cycloheximide were purchased from Sigma-Aldrich. Recombinant mouse TNF-α was from R&D systems (Minneapolis, MN). Thiamet-G (TMG) and z-VAD (OMe)-FMK were from Cayman Chemical (Ann Arbor, MI). Hydroxyproline assay kit (Sigma-Aldrich, St. Louis, MO) and ALT activity assay kit (Cayman Chemical, Ann Arbor, MI) were purchased from indicated sources and the assays were performed according to the manufacturer’s protocol.

### Statistical analyses

Results are presented as mean±S.E.M. The comparisons were carried out using two-tailed unpaired Student’s t-test or one-way ANOVA followed by Tukey-adjusted multiple comparisons. Data were plotted with GraphPad Prism. Statistical tests used are stated in the figure legends.

## Results

### Impaired O-GlcNAc signaling in humans and mice with liver injury

To examine the involvement of O-GlcNAc modification in liver disease, we investigated the changes in O-GlcNAc signaling in patients with liver cirrhosis. The expression of α smooth muscle actin (αSMA) was greatly increased in patients, indicating excessive extracellular matrix (ECM) deposition and liver injury in these patients (Fig 1A-B). Global O-GlcNAc levels were decreased in the livers from cirrhosis patients, which was associated with decreased OGT protein expression and increased OGA protein expression (Fig 1A-B). We also found a significant increase in the level of phosphorylated MLKL, indicating the augmentation of necroptosis in patients with liver cirrhosis. These results show that both O-GlcNAc and necroptosis signaling are associated with the development of liver cirrhosis in patients. To test whether O-GlcNAc signaling is also altered in chronic liver disease, we examined the changes in O-GlcNAc levels in a mouse model of ethanol-induced liver injury. Decreased liver O-GlcNAc levels were seen in mice fed with the ethanol diet as compared with the control group, which could be explained by the reduced expression of OGT in ethanol-fed mice (Figure 1C). It has been shown that necroptosis is induced in the mouse model of alcoholic liver disease (37). Taken together, these results indicate that the downregulation of O-GlcNAcylation is correlated with the induction of necroptosis in humans and mice with chronic liver injury.

**Figure 1.**
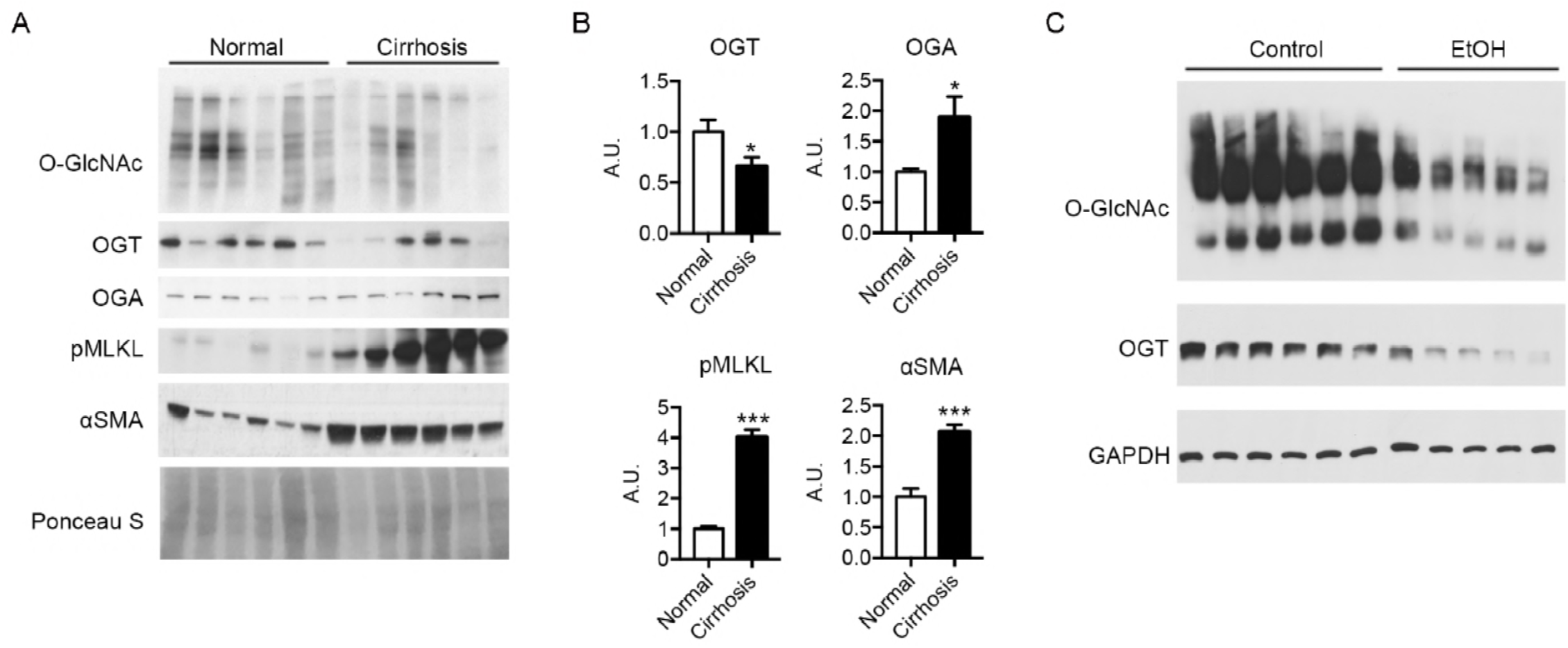
Defective O-GlcNAc signaling in patients with liver cirrhosis and in mice with ethanol-induced liver injury. (A) Western blots showing the protein expression in livers from healthy control and cirrhosis patients (n = 6). (B) Densitometric analysis of western blot results. A.U., arbitrary unit. (C) Western blots showing the protein expression in livers from control or ethanol diet-fed mice. n = 6 for control group and n = 5 for ethanol diet-fed group.

### Loss of OGT in hepatocytes leads to rapidly developed hepatomegaly and ballooning degeneration in mice

To investigate the physiological roles of OGT in the liver, we generated hepatocyte-specific OGT knockout mice (OGT-LKO) and control littermates (WT) by crossing *Albumin-Cre; Ogt^F/Y^* mice with *Ogt^F/F^* mice. Both mRNA and protein levels of OGT were significantly reduced in OGT-LKO mouse livers, confirming knockout efficiency (Figure 2A; Figure EV1A). As a result of OGT deletion, global O-GlcNAc levels were also diminished in hepatocytes as demonstrated by Western blot and immunohistochemistry (IHC) analyses (Figure 2A; Figure EV1B). WT and OGT-LKO mice had comparable body weight and showed no difference in oxygen consumption, food intake, and physical activities when measured at 4, 10, and 24-weeks of age (Figure EV2, data shown were measured at 10 weeks of age). However, at 4 weeks of age, OGT-LKO mice exhibited hepatomegaly and elevated circulating alanine aminotransferase (ALT) and aspartate aminotransferase (AST) levels, indicating a rapid development of liver injury in these mice (Figure 2B-E; Figure EV1C-D). This injury was not due to developmental defects since no abnormality was identified in 1-week-old OGT-LKO mice (Figure EV1E). We performed pathological staining and scoring to further analyze the changes in the knockout mice. Histological analysis of Hematoxylin and eosin (H&E)-stained liver sections identified ballooning degeneration in the OGT-LKO liver, as shown by swollen hepatocytes, vacuolated cytoplasm, and accumulation of Mallory hyaline (Figure 2F-G). Mild collagen deposition and sinusoidal fibrosis were observed in Masson’s trichrome stains (Figure 2H). Although the hydroxyproline content was not significantly higher in the OGT-LKO liver (Fig 2I), pathology scores revealed that 4-week-old knockout mice were in early stages of liver fibrosis (16.7% in stag1a, 66.7% in stage 1c, and 16.6% in stage 2) whereas all WT mice were healthy (Figure 2J). Alongside the histological observations, the mRNA levels of fibrogenic genes (*Acta-2, Col1a1,* and *Mmp2*) and pro-inflammatory genes (*Ccl2, Il-6,* and *Tnf-α)* were increased in 4-week-old OGT-LKO liver (Figure 2K-L). Pro-inflammatory cytokines, TNF-α and IL-6, were increased both in the liver and in the plasma of OGT-LKO mice (Figure 2M-N). Together, these results demonstrate that OGT-LKO mice develop spontaneous liver damage in 4 weeks.

**Figure 2.**
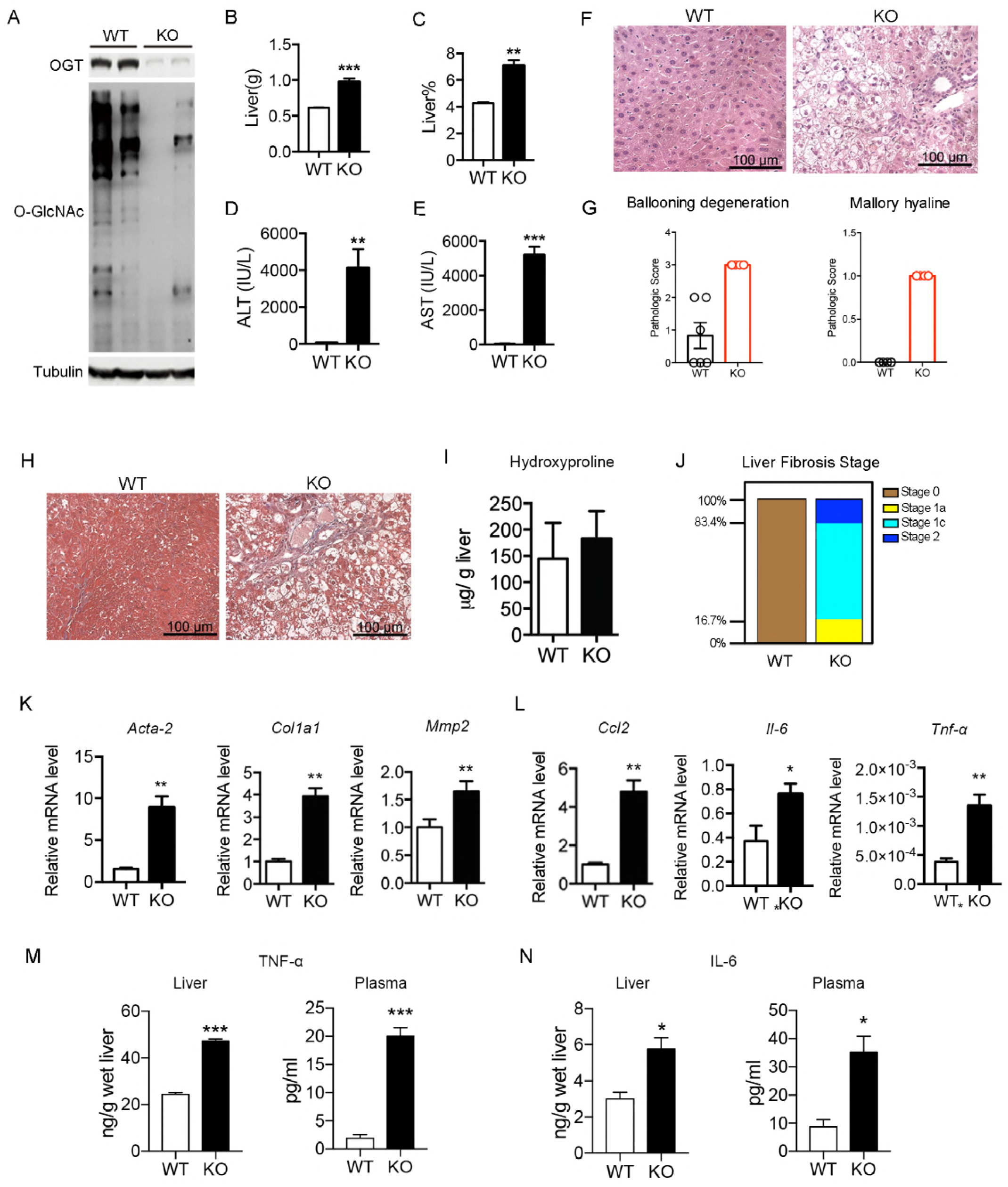
Hepatocyte-specific deletion of O-GlcNAc transferase (OGT) leads to hepatomegaly and liver injury in 4-week-old knockout mice. (A) Western blots showing the deletion of OGT and decrease of O-GlcNAc levels in liver-specific OGT knockout (OGT-LKO) mouse livers. (B) Liver weight. (C) Percentage of liver weight to body weight. (D-E) Alanine aminotransferase (ALT) and aspartate aminotransferase (AST) levels in the plasma from WT and OGT-LKO mice. (F-G) Hematoxylin and eosin (H&E) stains and pathologic scores of liver sections from WT and OGT-LKO mice. (H) Masson’s trichrome stains. (I) Hydroxyproline content. (J) Liver fibrosis stages in 4-week-old mice. (K) Expression of fibrogenic genes. (L) Expression of pro-inflammatory genes. (M-N) Liver and plasma TNF-α and IL-6 levels determined by ELISA (n = 3). n = 4, both genders. Data are shown as mean ± SEM. * *p* < 0.05; ** *p* < 0.01; *** *p* < 0.001 by unpaired Student’s t-test.

**Figure EV1.**
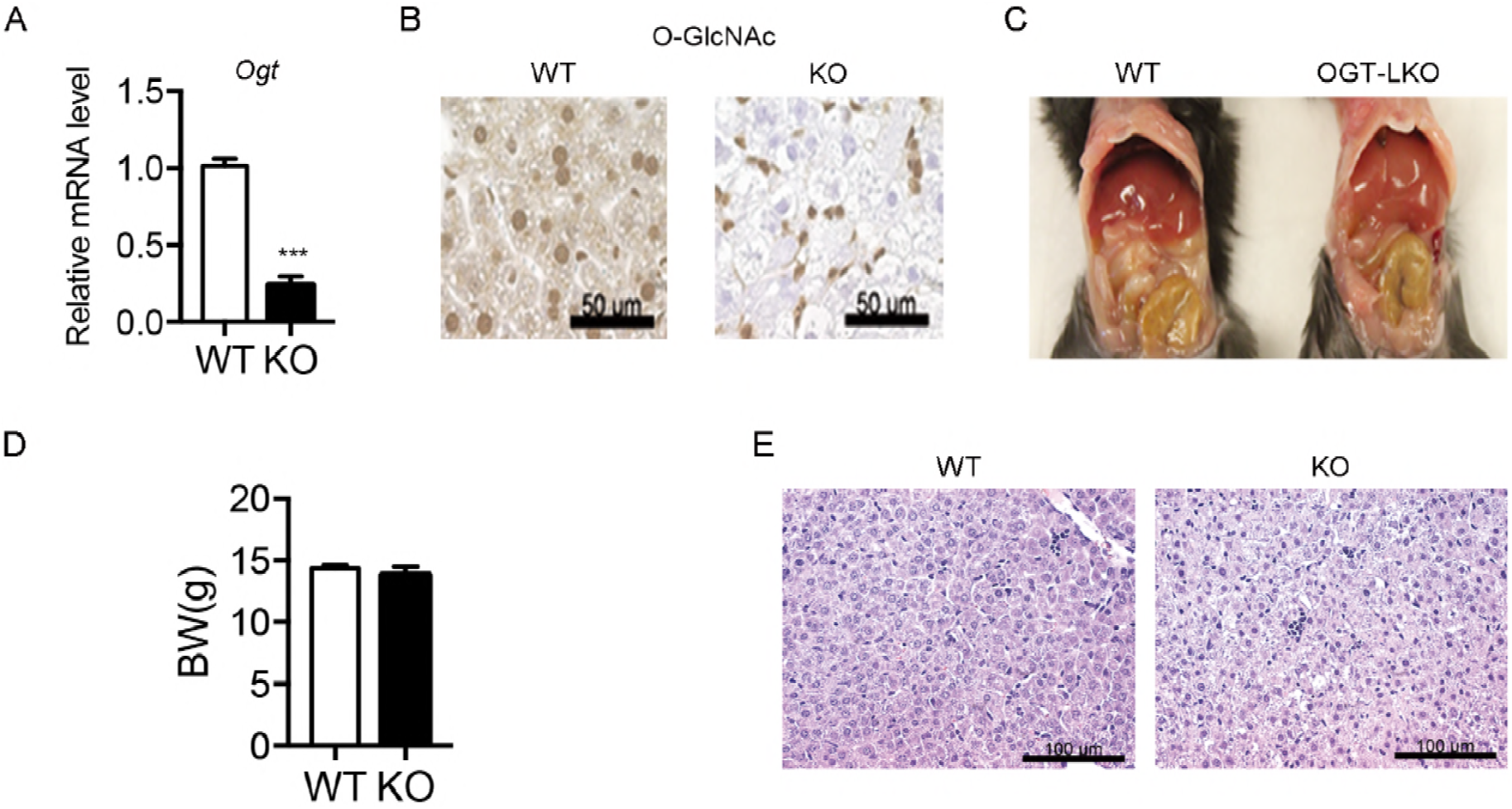
Deletion of OGT in the liver leads to hepatomegaly. (A) Gene expression of *Ogt* in WT and OGT-LKO livers. (B) IHC staining for O-GlcNAc in liver sections. (C) Morphology of livers from 4-week-old WT and OGT-LKO mice. (D) Body weight of 4-week-old WT and OGT-LKO mice. (E) H&E stains of 1-week-old WT and OGT-LKO mouse livers. Data are shown as mean ± SEM. *** *p* < 0.001 by unpaired Student’s t-test.

**Figure EV2.**
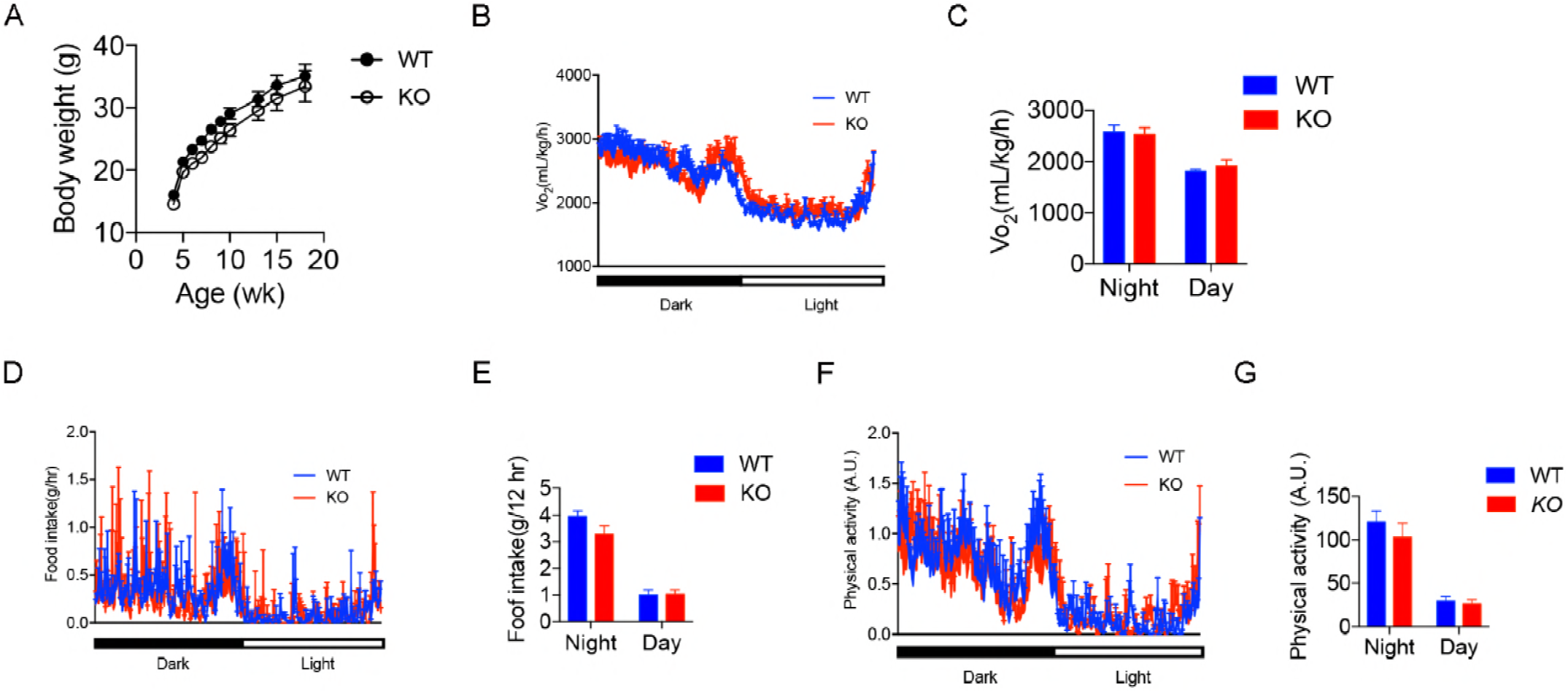
OGT-LKO mice show no metabolic abnormality. (A) Body weight monitoring of WT and OGT-LKO mice (n = 5). Real-time analysis of oxygen consumption rates (B), food intake (D), and physical activity (F) of WT and OGT-LKO mice. Average oxygen consumption rates (C), accumulative food intake (E), and accumulative physical activity (G) of WT and OGT-LKO mice. 10-week-old mice were used by the start of the metabolic cage studies. Data are shown as mean ± SEM.

### OGT deletion in hepatocytes leads to global transcriptome changes in the liver

Disruption of normal hepatocellular architecture and elevated expression of pro-inflammatory and fibrogenic genes indicate the possibility of progression to liver fibrosis, which is a critical phase in the development of chronic liver disease(38). Given the liver injury observed in OGT-LKO mice at 4 weeks, we performed RNA-sequencing to further analyze transcriptional changes in the OGT-LKO livers. The transcriptome analysis of the liver from 5-week-old OGT-LKO mice revealed profound changes in gene expression patterns as compared with WT mice (Figure 3A; Figure EV3A). 2341 genes showed at least 2-fold changes in expression between WT and OGT-LKO mice, among which 1525 genes were up-regulated and 816 genes were down-regulated (Figure 3B; Figure EV3B; Dataset S1). In agreement with our observation of early-onset fibrosis in 4-week-old OGT-LKO mice, gene set enrichment analysis revealed that the ECM – associated pathway was among the most enriched pathways in knockout livers (Figure 3C). Consistently, the mRNA expression levels of key mediators in ECM organization and collagen formation were elevated in OGT-LKO mice (Figure 3D). In contrast, genes that encode enzymes involved in amino acid or lipid metabolism, xenobiotic metabolism and detoxification were greatly repressed in OGT-LKO mice, indicating the impairment of major functions in hepatocytes from knockout mice (Figure 3C). Differentially expressed genes were also investigated with mouse phenotype enrichment analysis and gene-disease association analysis. Top results in these analyses were abnormal response to injury (MP: 0005164) and experimental liver cirrhosis (C0023893, DisGeNET). These results indicate that OGT-LKO mice quickly develop the fibrogenic program with global transcriptome changes and potentially serve as a genetic model to study liver fibrosis.

**Figure 3.**
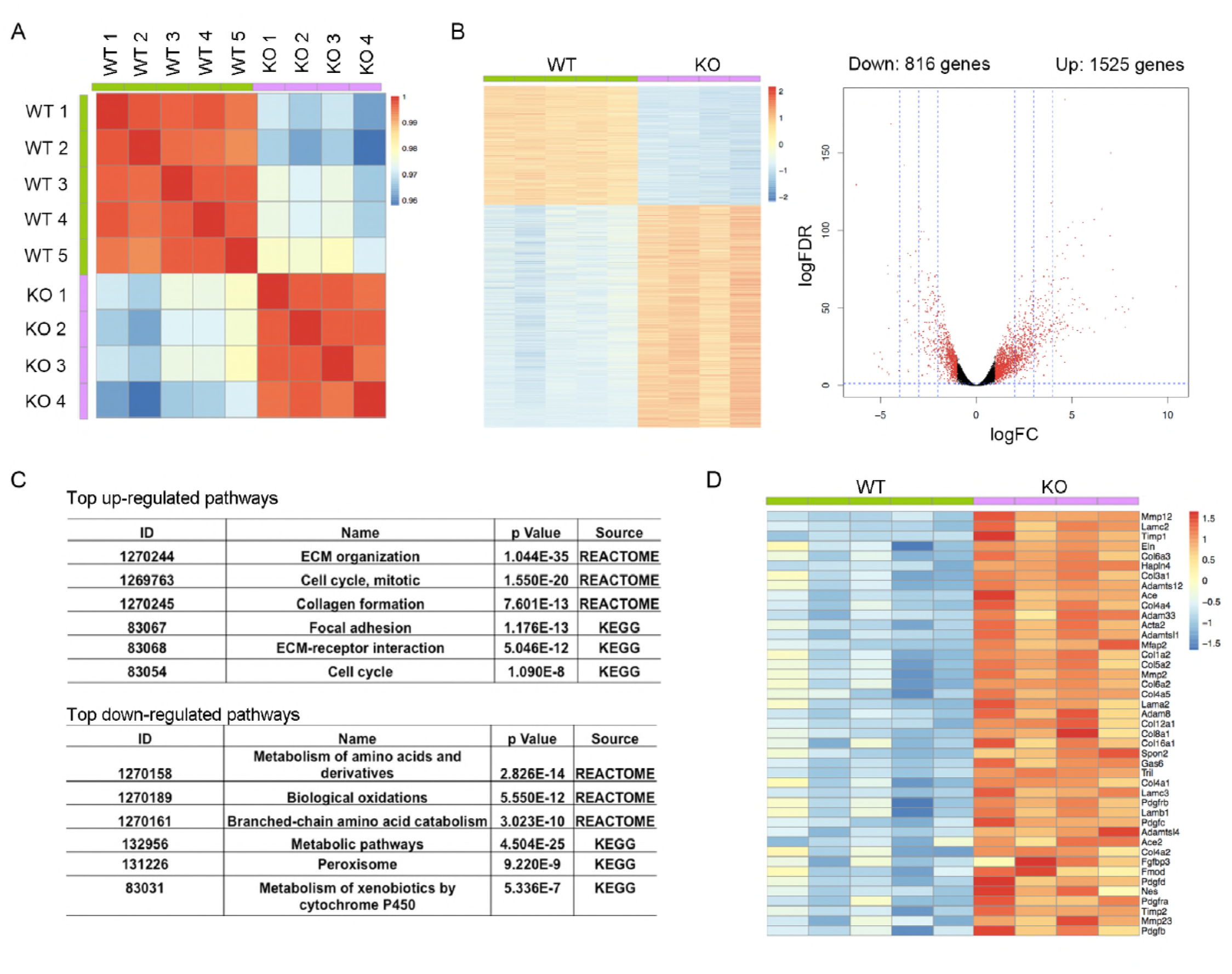
RNA-sequencing analysis reveals global transcriptional changes in OGT-LKO mouse livers at 5 weeks of age. (A) Heatmap of correlation matrix. (B) Heatmap of all differentially expressed genes with at least two-fold change in WT and OGT-LKO mice. These genes are in red in the volcano graph. FDR, false discovery rate. FC, fold change. (C) List of the top up-regulated (top) and down-regulated (lower) pathways in OGT-LKO mice analyzed with REACTOME and KEGG pathway enrichment analysis. (D) Representative genes in the extracellular matrix (ECM) organization pathway that are up-regulated in OGT-LKO livers.

**Figure EV3.**
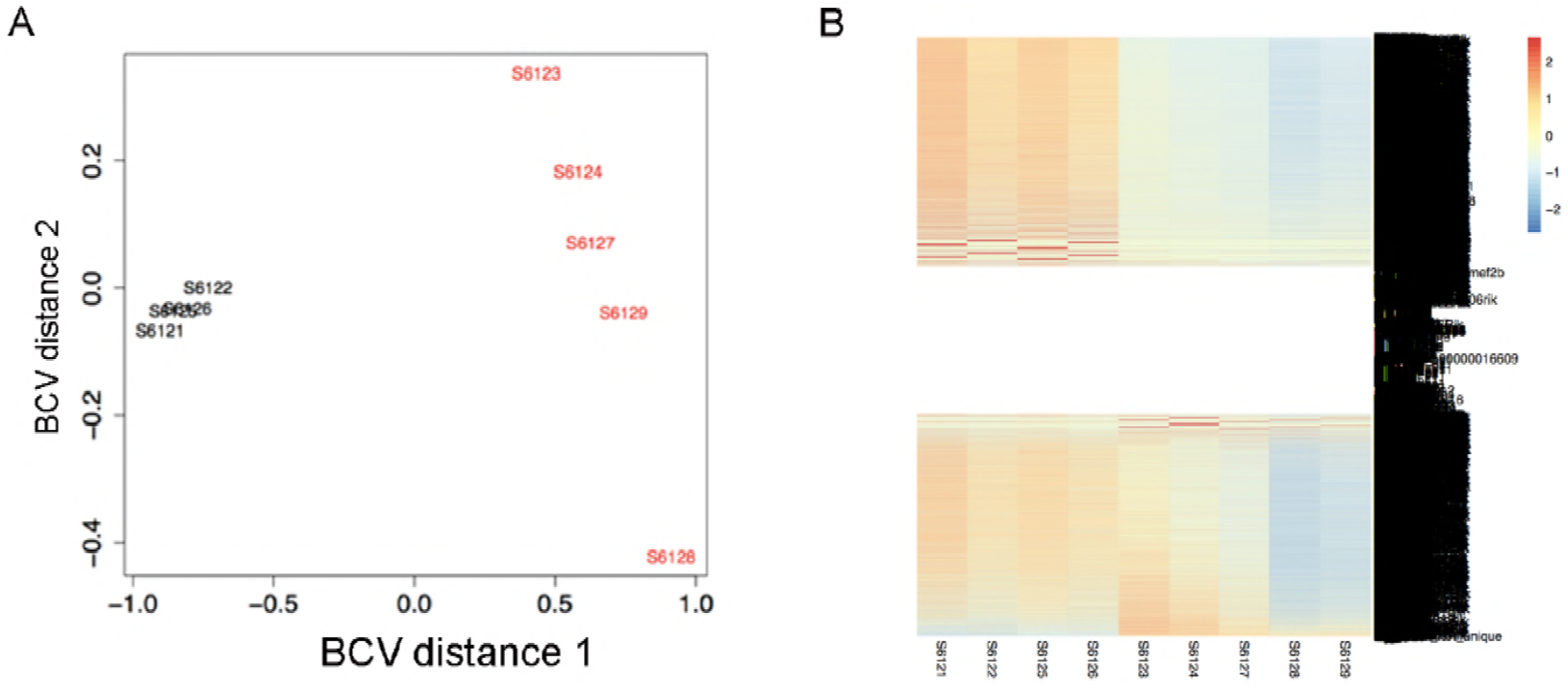
RNA-sequencing analysis of WT and OGT-LKO mouse livers at 5 weeks of age. (A) Multidimensional scaling (MDS) plot indicating the clustering of WT and KO samples. BCV, biological coefficient of variation. (B) Gene expression profile of all sequenced samples.

### OGT-LKO mice progress to liver fibrosis and portal inflammation at 10 weeks of age

We monitored the progression of liver injury in OGT-LKO mice and found that 10-week-old OGT-LKO mice exhibited more severe hepatomegaly with higher ALT and AST levels as compared with WT mice (Figure 4A; Figure EV4A-B). The H&E staining of OGT-LKO mouse liver sections showed massive immune cell infiltration that indicated portal and lobular inflammatory responses (Figure 4B). IHC analysis on F4/80 of these sections also showed an increase of infiltrating macrophages in the OGT-LKO livers (Figure EV4C). Excessive collagen deposition in the Masson’s trichrome stains revealed extensive sinusoidal fibrosis in OGT-LKO mice (Figure 4C). Pathological scoring of liver fibrosis, portal, and lobular inflammation confirmed the progression to liver fibrosis and inflammation in 10-week-old OGT-LKO mice (Figure 4D). This increase of infiltrating immune cells and ECM deposition also explain the enlargement of the livers in OGT-LKO mice. The spleen was also enlarged in OGT-LKO mice (Figure 4E), which could be partially attributed to portal hypertension caused by liver fibrosis in 10-week-old OGT-LKO mice. 10-week-old OGT-LKO livers also showed strikingly higher hydroxyproline content as compared with WT mice, which indicated elevated ECM production in the injured livers (Figure 4F). Consistent with the pathological observations of liver fibrosis and inflammation, OGT-LKO mice showed a dramatic increase in the expression of fibrogenic *(Acta-2, Col1a1,* and *Mmp2)* and pro-inflammatory genes *((Ccl2, Il-6,* and *Tnf-α)* (Figure 4G-H). Collectively, these results demonstrate that OGT-LKO mice showed a rapid and spontaneous progression from ballooning degeneration to liver fibrosis and inflammation within 10 weeks after birth.

**Figure 4.**
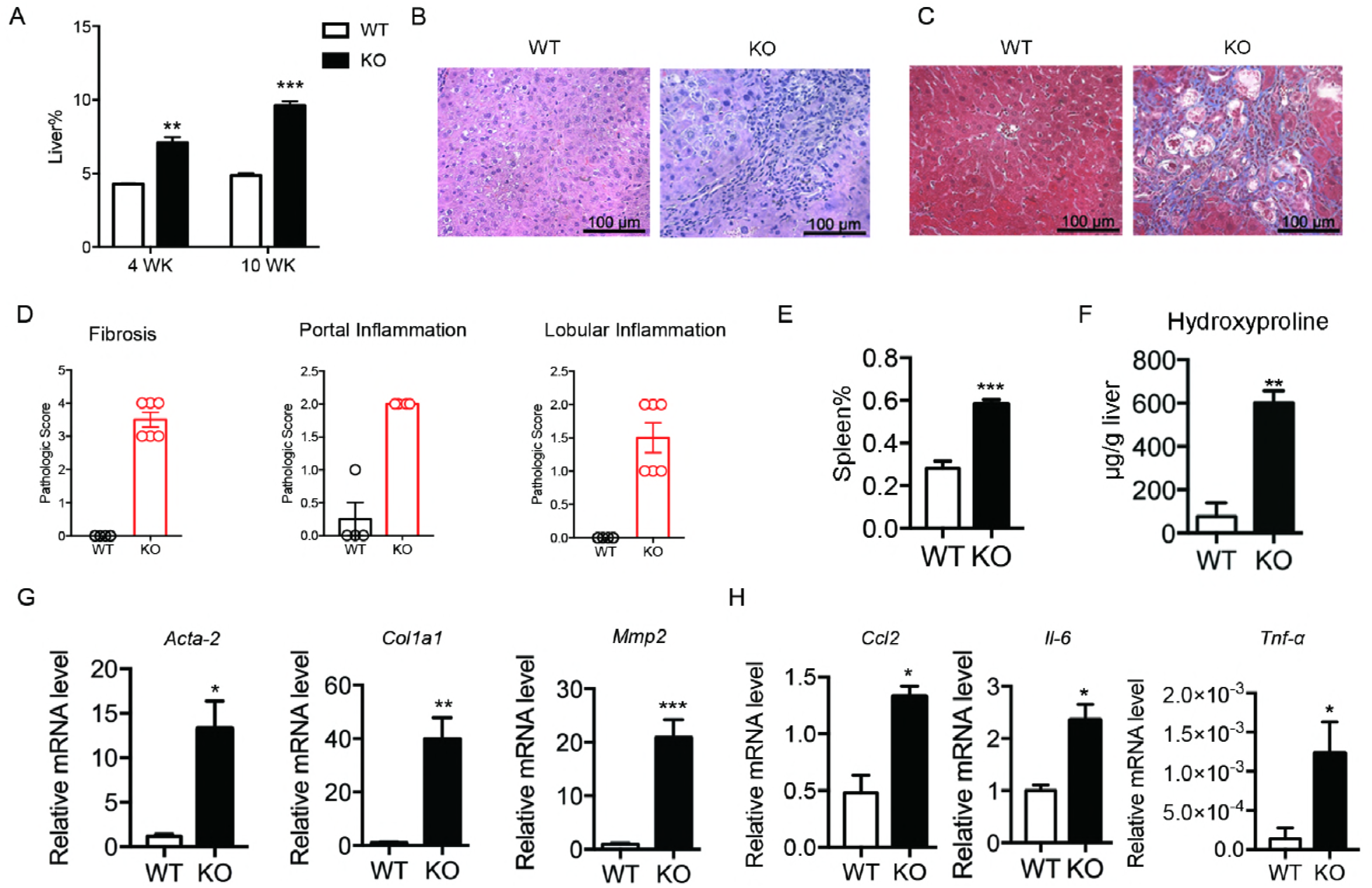
OGT-LKO mice develop liver fibrosis and inflammation at 10 weeks of age. (A) Percentage of liver weight to body weight in 4 week and 10 week-old mice. (B) H&E stains. (C) Masson’s trichrome stains. (D) Pathologic scores for liver fibrosis, portal and lobular inflammation. (E) Percentage of spleen weight to body weight. (F) Hydroxyproline content in the liver. (G) Expression of fibrogenic genes. (H) Expression of pro-inflammatory genes (n = 4). n = 6, both genders. Data are shown as mean ± SEM. * *p* < 0.05; ** *p* < 0.01; *** *p* < 0.001 by unpaired Student’s t-test.

**Figure EV4.**
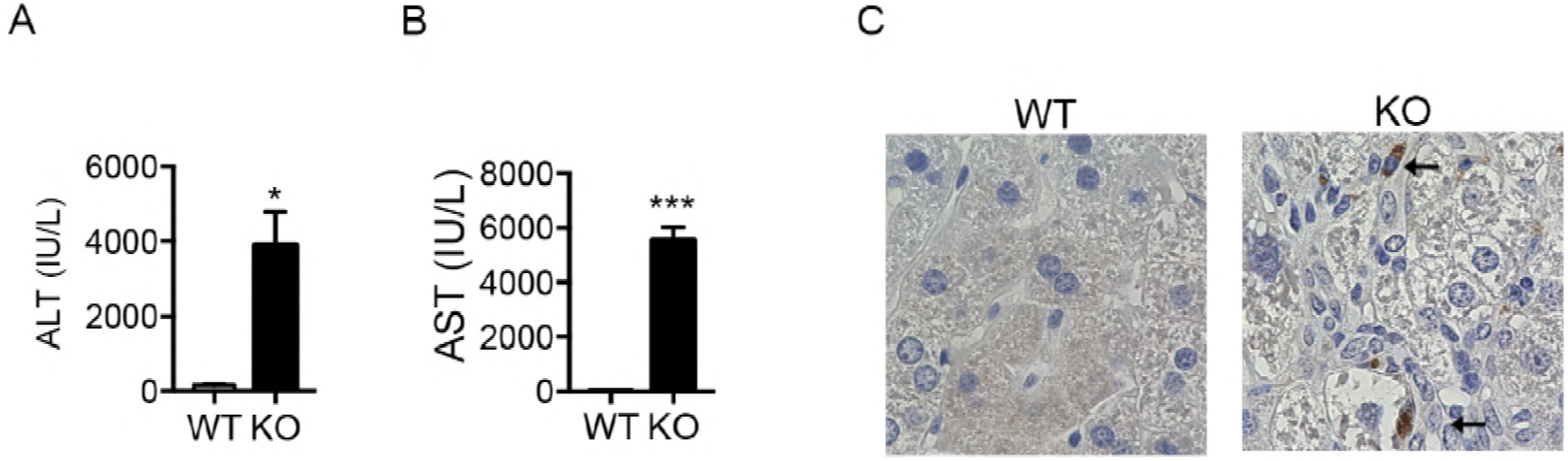
Exacerbated liver injury in 10-week-old OGT-LKO mice. (A) ALT levels. (B) AST levels. (C) IHC staining for macrophages in liver sections with F4/80 antibody. Arrows indicate representative positive stains. Data are shown as mean ± SEM. * *p* < 0.05; *** *p* < 0.001 by unpaired Student’s t-test.

### OGT-deficient primary hepatocytes undergo excessive necroptosis

To identify cellular dysfunction that leads to liver injury, we isolated primary hepatocytes from 4-week-old OGT-LKO and WT mice and plated the cells on collagen-coated plates. OGT-deficient hepatocytes exhibited rounded morphology and enlarged cell size (Figure 5A and EV5A). 24 h after plating, we examined the viability of isolated primary hepatocytes with Trypan blue staining and CellTiter assays. Concomitantly, viable cell number and cellular ATP content were both reduced in OGT-deficient hepatocytes, all of which represented necrotic cell death (Figure 5B-C). In primary hepatocytes stained with propidium iodine (PI), OGT-LKO hepatocytes also showed higher PI-positive population, indicating an increase in necrosis (Figure EV5B). To further determine which cell death pathway is involved in OGT-LKO hepatocellular death, primary hepatocytes were subjected to Annexin V-PI double staining and flow cytometry analysis. Compared to WT hepatocytes, a dramatic increase in the percentage of necroptotic cells was found in knockout hepatocytes whereas the apoptotic cell percentage was comparable between the two groups (Figure 5D). To confirm that apoptosis is not the major type of cell death in OGT-deficient hepatocytes, we examined the caspase activity in primary hepatocytes. Caspase-8 is a key mediator in the apoptosis pathway, activation of which initiates the caspase cascade and suppresses the necroptosis pathway. We found no changes in the activity of caspase-8 and its downstream targets caspase-3 and 7, indicating a minimal contribution of apoptosis to the cell death seen in OGT-deficient hepatocytes (Figure 5 E-F). The execution of necroptosis relies on the kinase activity of RIPK3, which could be inhibited by a selective inhibitor GSK-872. To test the hypothesis that OGT-LKO primary hepatocytes undergo massive necroptosis, we treated the cells with GSK-872 and found this necroptosis inhibitor significantly alleviated cell death in OGT-deficient hepatocytes (Figure 5G). In contrast, knockout hepatocytes treated with the pan-caspase inhibitor z-VAD showed no difference in cell viability as compared to the non-treated group, further confirming that apoptosis is not the major cause of hepatocyte death in OGT-LKO livers (Figure EV5C). Mitochondrial reactive oxygen species (ROS) production has been shown to be essential for necroptosis in many cell types (39). We therefore tested whether mitochondrial ROS production was increased in knockout hepatocytes. OGT-LKO hepatocytes displayed more intensive MitoSox staining as compared with WT hepatocytes (Figure 5H). Taken together, these results suggest that deficient O-GlcNAcylation induces necroptosis in hepatocytes.

**Figure 5.**
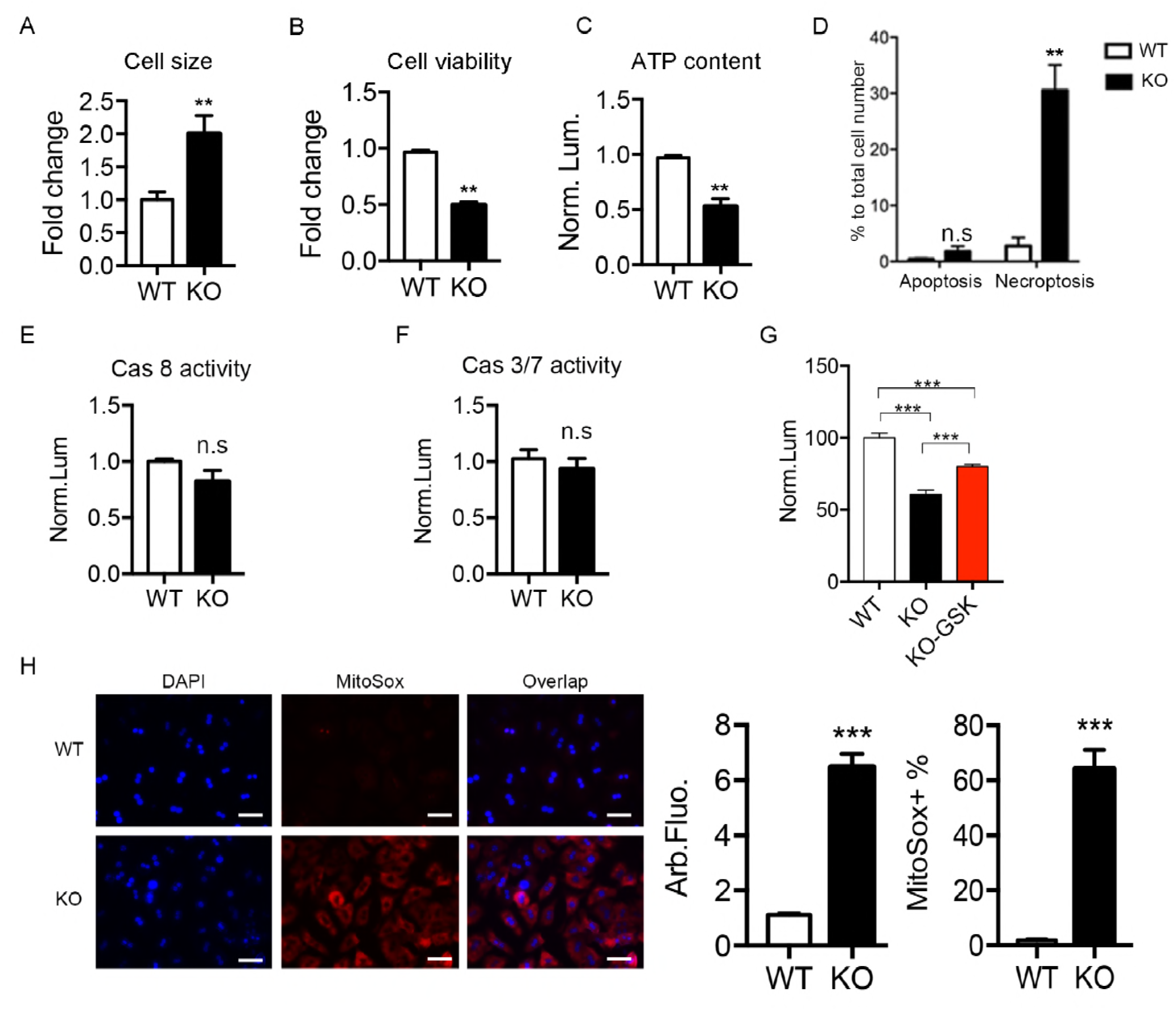
Loss of OGT in hepatocytes leads to excessive necroptosis. (A) Primary hepatocyte cell size quantified under light microscopy (n = 50). (B) Primary hepatocyte viability determined by Trypan blue staining (n = 3). (C) ATP content determined by the CellTiter-Glo luminescent cell viability assay kit (n = 3). Data were normalized to cell number. (D) Flow cytometry analysis of Annexin V-propidium iodine (PI) double-stained primary hepatocytes. Data were normalized to total cell number (n = 3). (E-F) Caspase-8 and caspase-3/7 activities in primary hepatocytes (n = 6). (G) Cell viability determined with the CellTiter-Glo kit. GSK, GSK-872, RIPK3 kinase inhibitor (n = 6). (H) MitoSox stains of primary hepatocytes. Fluorescence intensity and MitoSox-positive cell percentage were quantified with ImageJ (n = 20). Scale bar: 50 μm. Data are shown as mean ± SEM. * *p* < 0.05; ** *p* < 0.01; *** *p* < 0.001 by unpaired Student’s ř-test.

**Figure EV5.**
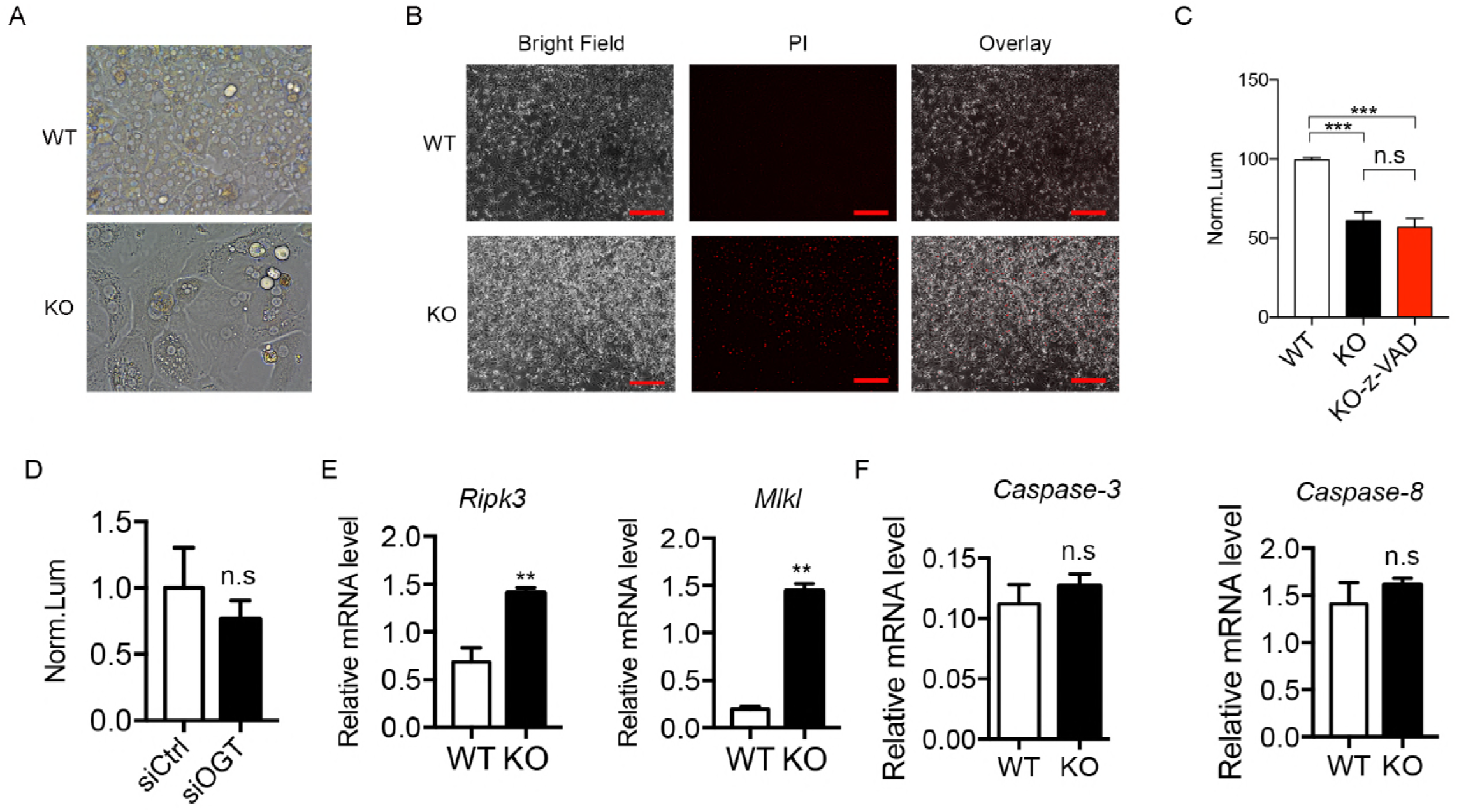
OGT-LKO hepatocytes show necroptotic, but not apoptotic, features. (A) WT and OGT-LKO primary hepatocytes examined with light microscopy. (B) PI staining for WT and OGT-LKO primary hepatocytes. Scale bar: 600 μm. (C) Cell viability determined with CellTiter-Glo kit. z-VAD is a pan-caspase inhibitor (n = 6). (D) WT primary hepatocytes were transfected with scrambled siRNA or OGT siRNA and treated with TNF-α and CHX for 24 hr to induce apoptosis. Cell viability was determined with CellTiter-Glo kit (n = 3). Data were normalized to the DMSO-treated group. (E) Gene expression of *Ripk3* and *Mlkl* in primary hepatocytes. (F) Gene expression of *Caspase-3* and *Caspase-8* in primary hepatocytes (n = 5). Data are shown as mean ± SEM. ** *p* < 0.01; *** *p* < 0.001 by unpaired Student’s t-test.

### OGT negatively regulates the necroptotic pathway

In multiple organs, increasing O-GlcNAcylation promotes cell survival (25). We hypothesized that OGT plays a pro-survival role in hepatocytes through suppressing necroptosis. To assess whether deletion of OGT selectively sensitized hepatocytes to necroptosis, but not apoptosis, primary hepatocytes from WT mice were transfected with scrambled siRNA or OGT siRNA. Apoptosis was induced by treating the cells with TNFα and cycloheximide (CHX) for 24 hr and OGT-ablated hepatocytes showed a similar response to the treatment as compared with the control group (Figure EV5D). The pan-caspase inhibitor z-VAD was added together with TNFα and CHX for 24 hr to induce hepatocyte necroptosis. Hepatocyte viability was significantly decreased in siOGT-transfected cells but not in the control group (Figure 6A). RIPK3 and MLKL are key mediators of necroptotic cell death and their phosphorylation indicates the activation of the necroptotic pathway(7). To investigate whether OGT deletion augments necroptosis by affecting RIPK3 and MLKL expression, we performed immunofluorescence staining of primary hepatocytes with antibodies against RIPK3 and MLKL. In agreement with our hypothesis, OGT-LKO hepatocytes showed increased immunofluorescence intensity of RIPK3 and MLKL staining as compared to WT hepatocytes (Figure 6B). Consistently, total and phosphorylated RIPK3 and MLKL proteins were increased in OGT-deficient hepatocytes whereas RIPK1, cleaved caspase-3, and cleaved caspase-8 protein expression levels were unchanged (Figure 6C), suggesting specific activation of necroptosis in OGT-LKO hepatocytes. IHC staining also revealed an increased expression of RIPK3 in OGT-LKO livers (Figure 6D). We also found an increase in *Ripk3* and *Mlkl* transcripts in OGT-LKO primary hepatocytes (Figure EV5E). In contrast, the mRNA expression levels of caspase-3 and caspase-8 were not changed in OGT-LKO hepatocytes (Figure EV5F). To test whether elevated RIPK3 and MLKL expression underlies the vulnerability to necroptotic treatments upon acute OGT deletion, primary hepatocytes transfected with scrambled siRNA or OGT siRNA were treated with TNFα, CHX, and z-VAD to induce necroptosis. Increased RIPK3 and MLKL protein levels were only seen in OGT siRNA-transfected cells (Figure 6E). Taken together, these data suggest that OGT negatively regulates necroptosis by repressing RIPK3 and MLKL expression.

**Figure 6.**
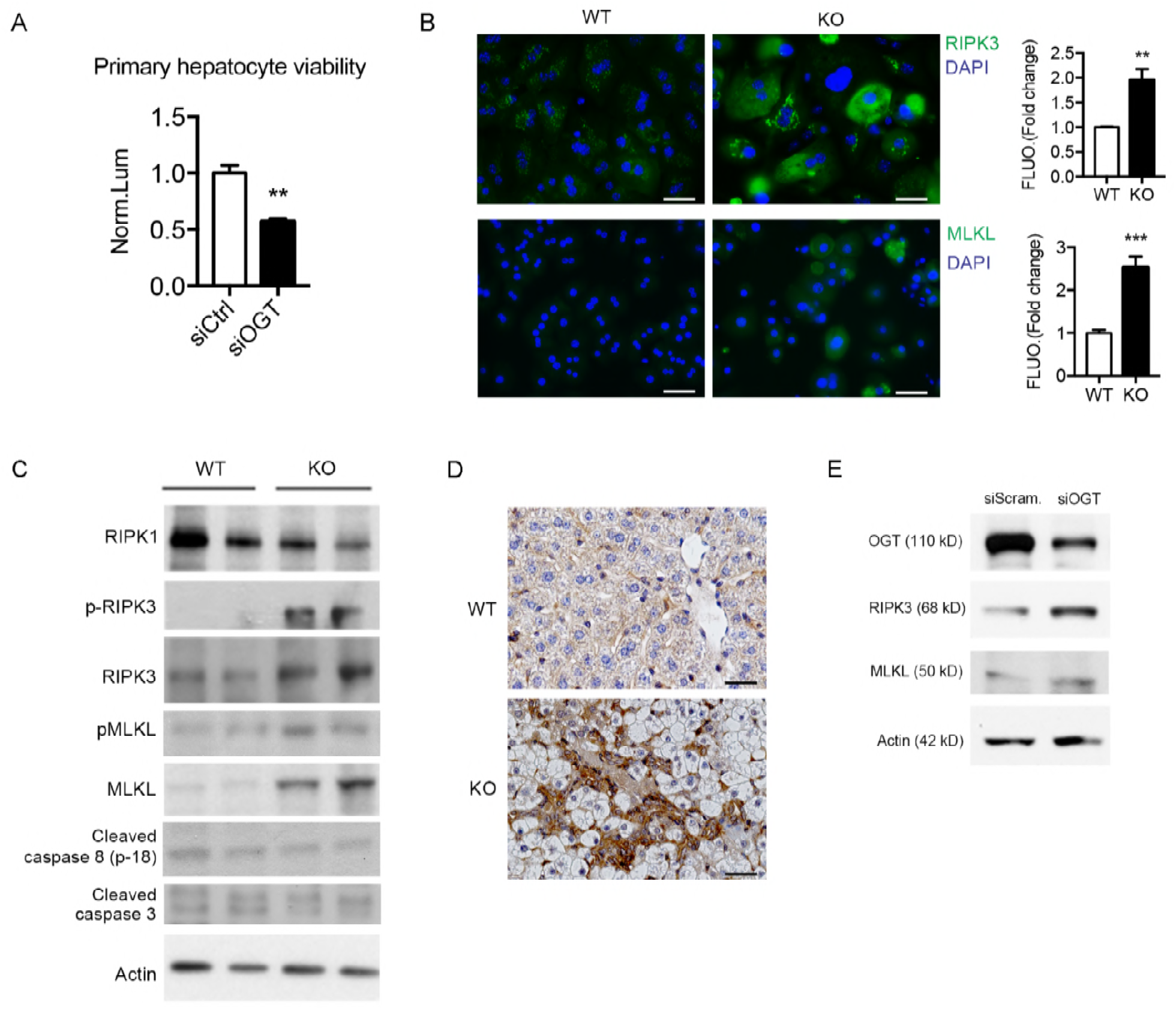
OGT deletion induces the expression of receptor-interacting protein kinase 3 (RIPK3) and mixed lineage kinase domain-like (MLKL) in primary hepatocytes. (A) WT primary hepatocytes transfected with scrambled siRNA or OGT siRNA were treated with TNF-α, cycloheximide (CHX), and z-VAD for 24 hr to induce necroptosis. Cell viability was determined with CellTiter-Glo luminescent cell viability assay kit (n= 3). Data were normalized to the DMSO-treated group. (B) Immunofluorescence images of primary hepatocytes stained with antibodies against RIPK3 and MLKL (green). The nucleus was stained with DAPI (blue). Fluorescence intensity was quantified with ImageJ (n = 30). (C) Western blotting of proteins from primary hepatocytes showing the expression levels of receptor-interacting protein kinase 1 (RIPK1), phosphorylated and total RIPK3, phosphorylated and total MLKL, cleaved caspase-8 and caspase-3. Actin was used as loading control. (D) Immunohistochemistry (IHC) stains of RIPK3 in liver sections from 4-week-old WT and OGT-LKO mice. (E) Western blotting of RIPK3 and MLKL in primary hepatocytes transfected with scrambled siRNA or OGT siRNA. Actin was used as loading control. All experiments were repeated at least three times with different cohorts of mice. Data are shown as mean ± SEM. * *p* < 0.05; ** *p* < 0.01; *** *p* < 0.001 by unpaired Student’s t-test.

### OGT glycosylates RIPK3 and decreases RIPK3 protein stability

We next sought to dissect the molecular mechanisms underlying OGT regulation of necroptosis. The balance of different PTMs, including O-GlcNAcylation, phosphorylation, and ubiquitination, precisely controls the expression and activity of their substrates. Previous studies have shown that RIPK3 is modified by both phosphorylation and ubiquitination(40, 41). We found abundant RIPK3 O-GlcNAcylation in WT mouse livers and HEK293T cells transfected with RIPK3 but to a much lower level in OGT-LKO mouse livers (Figure 7A). To determine whether OGT interacts with RIPK3, HEK293T cells were co-transfected with Myc-OGT and FLAG-RIPK3. Immunoprecipitation followed by Western blotting revealed the binding of OGT and RIPK3 (Figure 7B). To understand how OGT affects RIPK3 protein expression, increasing doses of the OGT-expressing plasmid were transfected into HEK293T cells. The result showed a dose-dependent decrease in RIPK3 proteins with increasing OGT expression (Figure 7C). To test whether glycosylation of RIPK3 affects its protein stability, a specific OGA inhibitor, Thiamet-G (TMG), was used to increase intracellular O-GlcNAc levels. Concomitantly, TMG effectively shortened the half-life of RIPK3 in H1299 cells as compared to the vehicle control (Figure 7D). Collectively, these results indicate that O-GlcNAcylation of RIPK3 decreases its protein stability and expression.

**Figure 7.**
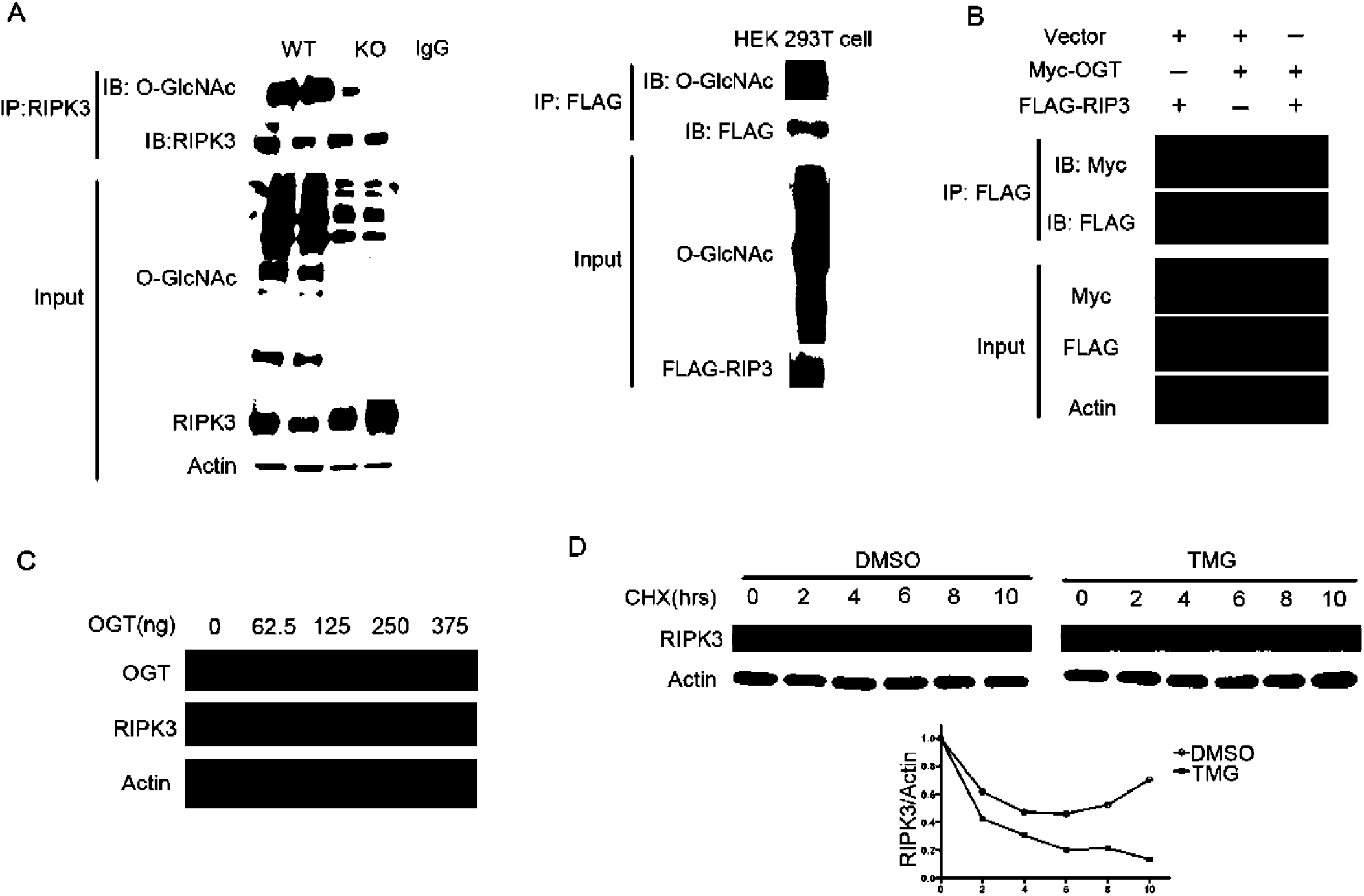
OGT glycosylates RIPK3 and regulates RIPK3 protein stability. (A) O-GlcNAcylation of RIPK3 in WT and OGT-LKO livers (left) and HEK293T cells transfected with RIPK3 (right) are shown. (B) HEK293T cells were co-transfected with Myc-OGT and FLAG-RIPK3, and their interaction was determined. Empty vector plasmid was transfected to keep equal transfection amount in each group. (C) HEK293T cells were co-transfected with RIPK3 and various doses of OGT as indicated. Levels of RIPK3 and OGT were determined by Western blots. (D) RIPK3-transfected H1299 cells were pre-treated with DMSO or TMG and then with CHX for indicated time points. Stability of RIPK3 was determined with Western blots and quantified with ImageJ. All experiments were repeated at least twice.

## Discussion

Liver fibrosis and cirrhosis is the common path to hepatocellular carcinoma (HCC), which is among the most deadly cancer around the world. Hepatocellular death exists in almost all acute and chronic liver pathologies, acting through different cell death pathways to initiate either phagocytosis, or compensatory proliferation, or inflammation and fibrosis. Here, we report that OGT-deficient hepatocytes undergo massive necroptosis, which leads to fast-developing liver injury manifested by hepatocyte ballooning, ALT elevation, inflammation, and liver fibrosis. Moreover, apoptosis is not evident in our model, indicating that necroptosis itself could contribute significantly to the pathogenesis of liver injury. It has only been in very recent years that we started to appreciate that besides apoptosis, necroptosis is also involved in the etiology of ASH, NASH, drug-induced liver injury, and viral infections(37, 42–46). We observed robust activation of necroptosis in liver cirrhosis patients, confirming the critical role of this cell death pathway in liver diseases. Compared to other forms of cell death, necroptosis represents a more “inflamed” mode, raising a higher possibility of tissue inflammation and subsequent fibrogenic events. This phenomenon is well exemplified in our model as we observed an early onset of liver fibrosis and inflammation in 4-week-old OGT-LKO mice. Part of this may result from the release of DAMPs in necroptotic hepatocytes and the amplification of inflammation responses. However, we did not observe any further progression from liver fibrosis to cirrhosis or HCC in elder OGT-LKO mice, underscoring the significant contribution of necroptosis to fibrogenic liver pathologies(1).

In our study, we showed that the expression of RIPK3 is greatly induced in OGT-LKO primary hepatocytes, whereas the expression and activities of caspases are unchanged in these cells as compared with WT primary hepatocytes. Previous studies have proposed that an intricate balance and mutual regulation between apoptosis and necroptosis help maintain tissue homeostasis, and suppressing either of them will cause the up-regulation in the other(47, 48). The molecular basis for this is that the necroptosis pathway overlaps with the apoptosis pathway regarding the protein complex assembled upon death receptor binding. However, the two pathways diverge downstream of RIPK1 where either caspase-8 or RIPK3 is activated. In the liver, studies have shown that the reciprocal inhibition of caspase-8 and RIPK3 is critical for the development of NASH and hepatocarcinogenesis(6, 49). RIPK3 inhibits the cleavage caspase-8 and subsequent activation of JNK to stall cell proliferation and thus limit HCC development(49). Interestingly, a previous study by Xu.et al demonstrated that overexpression of OGT in the liver promotes HCC through activation of JNK(50). These results along with ours indicate that OGT is pro-survival in the hepatocytes, ablation of which not only results in uncontrolled necroptosis but may also compromise cell proliferation. Further investigations into the role of OGT in other chronic liver injury models would elucidate how OGT is integrated in this signaling network.

We have demonstrated that O-GlcNAcylation reduces RIPK3 protein stability and suppresses its expression. It has long been known that PTMs such as glycosylation, phosphorylation, and ubiquitination, add a critical layer of regulation on protein functions. In regard to the necroptotic pathway, phosphorylation of RIPK3 is best studied because it initiates the essential steps in necroptosis by recruiting and phosphorylating MLKL(51).

Previous studies have discovered that a great number of substrates could be modified by OGT or protein kinases at the same or proximal sites. We therefore hypothesize that RIPK3 is also a substrate for both glycosylation and phosphorylation and the competitive interplay between them controls the activity of RIPK3. In agreement with this hypothesis, several sites (including T248, T368, T374, T378, S379, T463, S466, and T467) have been predicted with the online PTM prediction tool (http://www.cbs.dtu.dk/services/YinOYang/). Mapping these specific regulatory sites will be an important subject for future investigation to better understand the molecular mechanisms governing the necroptotic pathway.

Liver fibrosis is a critical stage in chronic liver disease and very limited antifibrotic treatments are currently available. One of the difficulties in developing anti-fibrotic drugs is the lack of effective animal models. Available genetic models of liver fibrosis are mostly whole body knockout mice, such as *Mdr2*^-/-^(52), II-*2R*^-/-^(53), or dominant negative Tgf-βRII mice(54), which have confounding effects ascribed to other tissues. The OGT-LKO mouse model we report here can potentially serve as a novel, liver-specific genetic model for future research on liver fibrosis. Moreover, a majority of current mouse models are established with a special diet or toxic reagents. It takes at least several months to develop liver fibrosis and the phenotypes are usually irreversible. Therefore, many features in human liver fibrosis cannot be faithfully recapitulated and large-scale, long-term studies are often impossible in these models. In contrast, OGT-LKO mice spontaneously develop liver fibrosis in less than 10 weeks. The OGT-LKO model can thus be employed as a novel, effective mouse model of liver fibrosis with broad translational implications for anti-fibrotic drug screening and evaluation.

## Acknowledgements

We thank Dr. Xiaochun Yu from the University of Michigan for providing the Myc-OGT plasmid and Dr. Vishva Dixit at Genentech for providing the p-RIPK3 and MLKL antibodies. We thank Kathy Harry, Dr. Masatake Tanaka, Dr. Jittima Weerachayaphorn, and Dr. Mateus T. Guerra from Yale University Liver Center for their generous help with experimental procedures.

## List of Abbreviations

OGT: O-GlcNAc transferase
O-GlcNAc: O-linked β-N-acetylglucosamine
OGA: O-GlcNAcase
ALT: alanine aminotransferase
RIPK3: receptor-interacting protein kinase 3
MLKL: mixed lineage kinase domain-like
RIPK1: receptor-interacting protein kinase 1
ASH: alcoholic steatohepatitis
NASH: non-alcoholic steatohepatitis
HCC: hepatocellular carcinoma
DAMPs: damage-associated molecular patterns
PTM: post-translational modification
WT: wild type
OGT-LKO: liver-specific OGT knockout
H&E: hematoxylin and eosin
IHC: immunohistochemistry
ELISA: enzyme linked immunosorbent assays
RT-qPCR: real time-quantitative polymerase chain reaction
TNF-α: tumor necrosis factor alpha
IL-6: interleukin 6
PI: propidium iodine
TMG: thiamet G
DMSO: dimethyl sulfoxide
Acta-2: alpha actin 2
Col1a1: collagen type 1 alpha 1
Mmp2: metalloproteinase-2
Tgf-β: transforming growth factor beta
Ccl2: C-C motif chemokine ligand 2
ECM: extracellular matrix
CHX: cycloheximide
JNK: c-Jun N-terminal kinases
Sp1: specificity protein 1.

## Notes

**Financial support:** This work was supported by the National Institutes of Health (R01DK102648), the Yale Liver Center (P30 DK34989), and the American Cancer Society (RSG-14-244-01-TBE) to X.Y., the National Institutes of Health (P01-DK57751) to X.Y., A.M.B., B.E.E., M.E.R., and M.H.N.,, the China Scholarship Council-Yale World Scholars fellowship to B.Z..

